# Ageing disrupts MANF-mediated immune modulation during skeletal muscle regeneration

**DOI:** 10.1101/2022.07.20.500588

**Authors:** Neuza S. Sousa, Margarida F. Brás, Inês B. Antunes, Päivi Lindholm, Joana Neves, Pedro Sousa-Victor

## Abstract

Age-related decline in skeletal muscle regenerative capacity is multifactorial, yet, the contribution of immune dysfunction to regenerative failure is unknown. Here, we uncover a new mechanism of immune modulation operating during skeletal muscle regeneration that is disrupted in aged animals and relies on the regulation of macrophage function. The immune modulator MANF is induced following muscle injury in young mice, but not in aged animals, and its expression is essential for regenerative success. Regenerative impairments in the aged muscle are associated with defects in the repair-associated myeloid response similar to those found in MANF-deficient models and could be improved through MANF delivery. We propose that restoring the myeloid response through immune modulation is a promising therapeutic strategy to improve the regenerative capacity of aged muscles.

## Introduction

Age-related decline in regenerative capacity is the synergistic result of cell intrinsic impairments in somatic stem cells and alterations in the local and systemic environment coordinating the repair process^1,2^. The reversible nature of some of the age-associated changes affecting the regenerative environment has been demonstrated through heterochronic parabiosis experiments, where old tissue is exposed to a youthful circulatory environment, highlighting the potential of interventions in the aged tissue milieu to reverse regenerative decline^3,4^. The myeloid system is a potential candidate to mediate these rejuvenating effects^5-7^.

The skeletal muscle is a paradigmatic model to study age-related loss of regenerative capacity. Skeletal muscle regeneration is sustained throughout life by a population of adult muscle stem cells (MuSCs) and relies on a highly coordinated sequence of events, engaging several niche populations^8-10^. Immune cells infiltrate the skeletal muscle soon after injury and are responsible for essential functions, including the clearance of tissue debris and the coordinated regulation of MuSC function and other niche components^11,12^. Macrophages are the most abundant type of immune cells participating in muscle repair and originate from infiltrating monocytes that differentiate in situ into pro-inflammatory macrophages^12^. Regenerative success depends on a timely regulated phenotypic transition of these pro-inflammatory macrophages into pro-repair macrophages^12-14^.

Recently, we identified a novel immune modulatory function for Mesencephalic Astrocyte-derived Neurotrophic Factor (MANF)^15^, an endoplasmic reticulum (ER)-stress-inducible protein with pleiotropic effects in multiple organs^16-19^. We found that MANF is a systemic regulator of inflammation and tissue homeostasis during ageing, and one of the factors required in young blood to promote part of the rejuvenation effects elicited by heterochronic parabiosis^20^. Circulatory MANF levels decline with age in mice and humans, and MANF supplementation in old mice is sufficient to limit inflammation and tissue damage in the liver^20^ and preserve retinal homeostasis^21^. However, how a decline in MANF signalling affects the age-related loss of regenerative capacity remains unexplored.

## Results

### MANF is essential for muscle regeneration

To explore the involvement of MANF in muscle regeneration, we evaluated MANF expression following muscle injury. We observed a sharp increase in MANF levels that peaked at 3-4 days post injury (dpi) and progressively declined thereafter (Fig. 1a-b and Extended data 1a). Importantly, this injury-dependent induction of MANF was blunted in aged animals (Fig. 1c and Extended data 1b), revealing an inability of aged muscles to significantly induce MANF between 2 and 3dpi (Extended data 1c).

**Figure 1.**
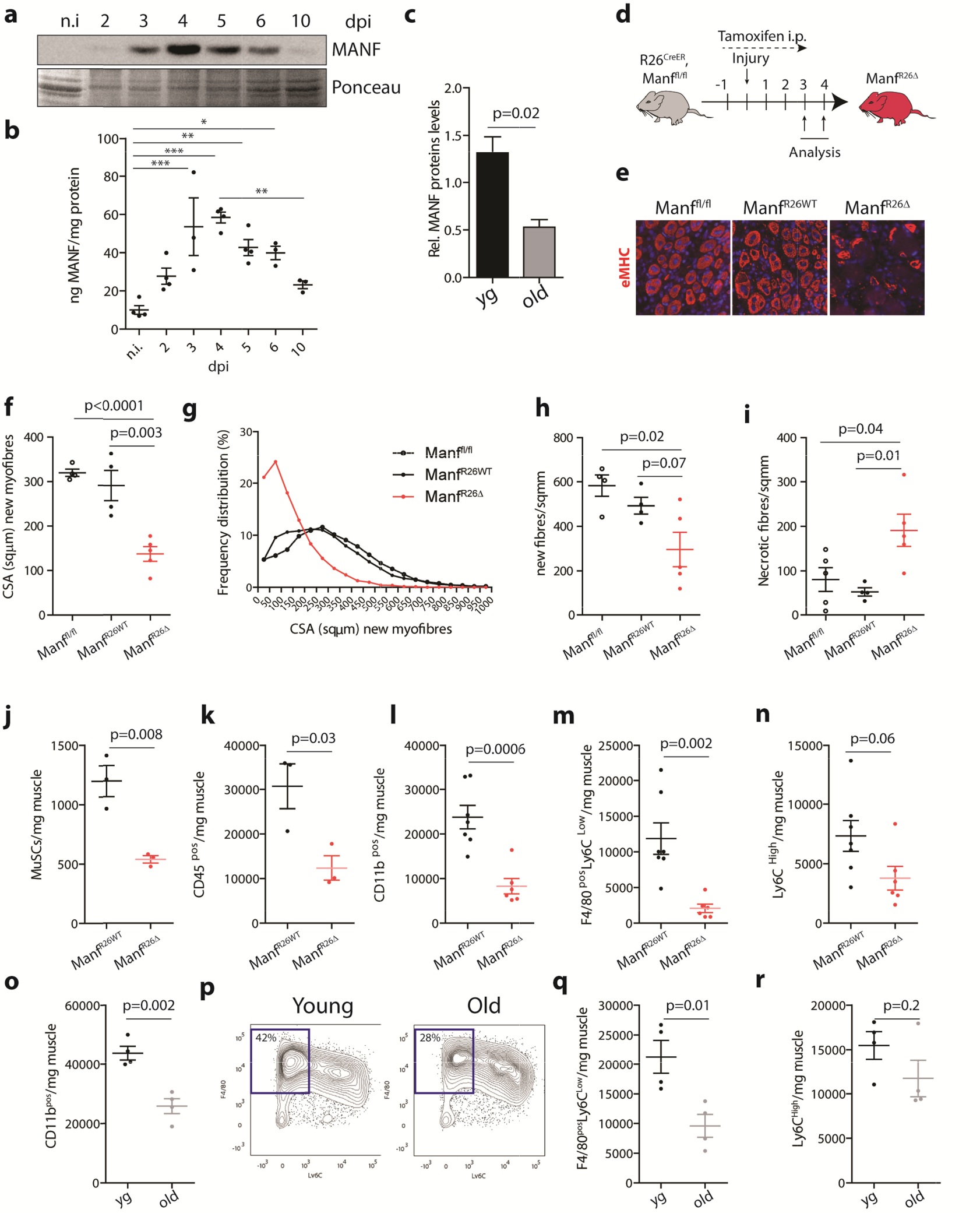
MANF is essential for skeletal muscle regeneration. **a**, Western blot analysis of MANF levels in protein extracts of tibialis anterior (TA) muscles from yg wild type (wt, C5 BL/6) mice, non-injured and at different time points following injury (2, 3, 4, 5, 6, 10 dpi). Ponceau S-staining of the membrane was used to verify equal protein loading in each sample. **b**, MANF protein levels, quantified by ELISA, in extracts of TA muscles from yg wt (C57BL/6) mice at different time points following injury (n=3-4/timepoint). **c**, Quantification of relative average levels of MANF, normalized to Ponceau levels, in protein extracts of TA muscles of yg (2-6mo) and old (22-25mo) wt (C57BL/6) mice at 3dpi (n=3-6/age, see Extended data Fig. 1b for representative Western blot images). **d**, Experimental timeline for analysis of animals with inducible and ubiquitous ablation of MANF. **e**, Representative images of eMHC (red) staining in cryosections of regenerating TA muscles from Manf^fl/fl^, Manf^R26WT^ and Manf^R26Δ^ mice, at 4dpi. DAPI is used to identify nuclei. Quantifications of this staining, for independent animals, are shown in Fig. 1f-h. **f-h**, Quantification of the average cross-sectional area of eMHC+ new myofibres (f), frequency distribution of new myofibres by size (g) and density of new myofibres (h) in regenerating TA muscles from Manf^fl/fl^, Manf^R26WT^ and Manf^R26Δ^ mice at 4dpi (n=4-5/condition). **i**, Quantification of the number of necrotic fibres in regenerating TA muscles from Manf^fl/fl^, Manf^R26WT^ and Manf^R26Δ^ mice at 4dpi (n=4-5/condition, see Extended data Fig. 1e for representative images). **j-n**, Quantification, by flow cytometry, of MuSCs (j), CD45^pos^ immune cells (k), myeloid cells (CD11b^pos^, l), pro-repair macrophages (F4/80^pos^Ly6C^Low^, m) and pro-inflammatory macrophages (Ly6C^High^, n) in regenerating quadriceps (QC) muscles of Manf^R26WT^ and Manf^R26Δ^ mice at 3dpi (j-k, n=3/condition; l-n, n=6-7/condition). **o**,**q**,**r**, Quantification, by flow cytometry, of myeloid cells (CD11b^pos^, o), pro-repair macrophages (F4/80^pos^Ly6C^Low^, q) and pro-inflammatory macrophages (Ly6C^High^, r) in regenerating QC muscles of yg (2-6mo) and old (22-24mo) wt (C57BL/6) mice at 3dpi (n=4/age). **p**, Representative density plots from flow cytometry analysis of muscle cell populations from yg and old animals based on F4/80 and Ly6C cell surface markers, at 3dpi. Percent numbers indicate pro-repair macrophages (F4/80^pos^Ly6C^Low^) relative to the CD11b^pos^ total population. Data are represented as average ± s.e.m and each n represents one animal. In b, p values are from one-way ANOVA with Bonferroni’s multiple comparison post-test. In all other graphs, p values are from two-tailed Student’s t-test. yg, young; n.i., non-injured; dpi, days post-injury; i.p., intraperitoneal; eMHC; embryonic myosin heavy chain; CSA, cross-sectional area; MuSCs, muscle stem cells.

To understand the consequences of defective MANF signalling for muscle regeneration, we generated a mouse model allowing the inducible and ubiquitous ablation of MANF in adult animals. In the Rosa26-Cre^ER^, Manf^fl/fl^ mice tamoxifen treatment immediately before muscle injury and during muscle regeneration (Fig. 1d) resulted in a complete loss of MANF protein in the regenerating tissue (Extended data 1d). Analysis of tamoxifen-treated Rosa26-Cre^ER^, Manf^fl/fl^ mice (herein referred to as Manf^R26Δ^), revealed impairments in muscle regeneration compared to oil treated mice (Manf^R26WT^) and tamoxifen-treated Manf^fl/fl^ mice. This is evidenced by a reduction in the density and the cross-sectional area (CSA) of new myofibres formed at 4dpi (Fig. 1e-h) and a persistence of necrotic myofibres within the regenerating muscle (Fig. 1i and Extended data 1e), resembling the phenotype found in aged animals^22,23^.

Manf^R26Δ^ mice also displayed a significant reduction in the number of MuSCs (p<0.01, Fig. 1j) and CD45^pos^ immune cells (p<0.05, Fig. 1k) in the regenerating muscle at 3dpi, with no changes in the populations of fibroadipogenic progenitors (FAPs) or endothelial cells (Extended data 1f-h). Considering the fundamental role of myeloid cells in the clearance of necrotic debris, a process impaired in Manf^R26Δ^ mice, we focused our analysis on the repair-associated myeloid response. Manf^R26Δ^ mice had a 65% reduction in the number of myeloid cells (CD11b^pos^) at 3dpi, corresponding to an 82% reduction in the pro-repair (F4/80^pos^Ly6C^Low^) macrophage population and a less pronounced reduction in the pro-inflammatory (Ly6C^High^) population (p=0.06, Fig. 1l-n and Extended data 1i). The neutrophil population was unchanged (Extended data 1j). Thus, MANF loss is associated with defects in the repair-associated myeloid response characterized by a reduced presence of myeloid cells in the injured muscle and an imbalance in macrophage states, whereby pro-inflammatory macrophages tend to accumulate at the expense of pro-repair macrophages.

### Ageing impairs the repair-associated myeloid response

Skeletal muscle from aged animals exhibits regenerative defects similar to those found in Manf^R26Δ^ mice^22,23^ and a blunted induction of MANF following injury (Fig. 1c and Extended data 1b-c). Thus, we asked whether this reduction in MANF levels in aged muscles was associated with similar defects in the repair-associated myeloid response. We found that 23-25mo old animals had a 41% reduction in the number of myeloid cells within the regenerating skeletal muscle at 3dpi (Fig. 1o and Extended data 1k), with a 55% reduction in the number of pro-repair macrophages and no significant changes in the number of pro-inflammatory macrophages (Fig. 1p-r). Importantly, these defects manifested as a reduced capacity to increase the total number of myeloid cells and pro-repair macrophages between 2 and 3 dpi (Extended data 1l-n). Thus, aged muscles display defects in myeloid cell accumulation and an imbalance of macrophage subpopulations that can be recapitulated in conditions of MANF loss. This parallel in myeloid defects included only the macrophage populations and not the neutrophil population, which was reduced in aged animals but not in Manf^R26Δ^ mice (Extended data 1j-k).

### MANF is specifically expressed in pro-repair macrophages during muscle regeneration

Since MANF is induced in macrophages after injury in other systems^15^ and macrophage numbers increase in the skeletal muscle following injury (Extended data 2a, see also REF^12^), we hypothesized that macrophages could be a source of MANF in the regenerating skeletal muscle. Indeed, F4/80^pos^ cells immunostained in muscle cryosections co-localized with sites of highest MANF expression (Extended data 2b). To further test this hypothesis we treated mice with clodronate liposomes during muscle injury, generating a condition where macrophage numbers are reduced by 80% but neutrophils are not affected (Fig. 2a and Extended data 2c), and observed a significant reduction in the levels of MANF protein present in the skeletal muscle at 3dpi (p<0.05, Fig. 2b). To distinguish whether the increase in MANF levels is due to the influx of macrophages following muscle injury or the induction of MANF within the macrophage population, we isolated F4/80^pos^ cells by fluorescence activated cell sorting (FACS) and analyzed MANF protein expression. Interestingly, MANF protein levels were also changed in the F4/80^pos^ population of macrophages (Fig. 2c), mimicking the expression dynamic observed in whole muscles (Fig. 1a-b) and following the phenotypic transition of pro-inflammatory macrophages into pro-repair macrophages during muscle repair (Extended data 2d, see also REF^24^). Consistently, analysis of isolated macrophage subpopulations revealed that MANF is specifically expressed in the F4/80^pos^Ly6C^Low^ subpopulation of pro-repair macrophages (Fig. 2d).

**Figure 2.**
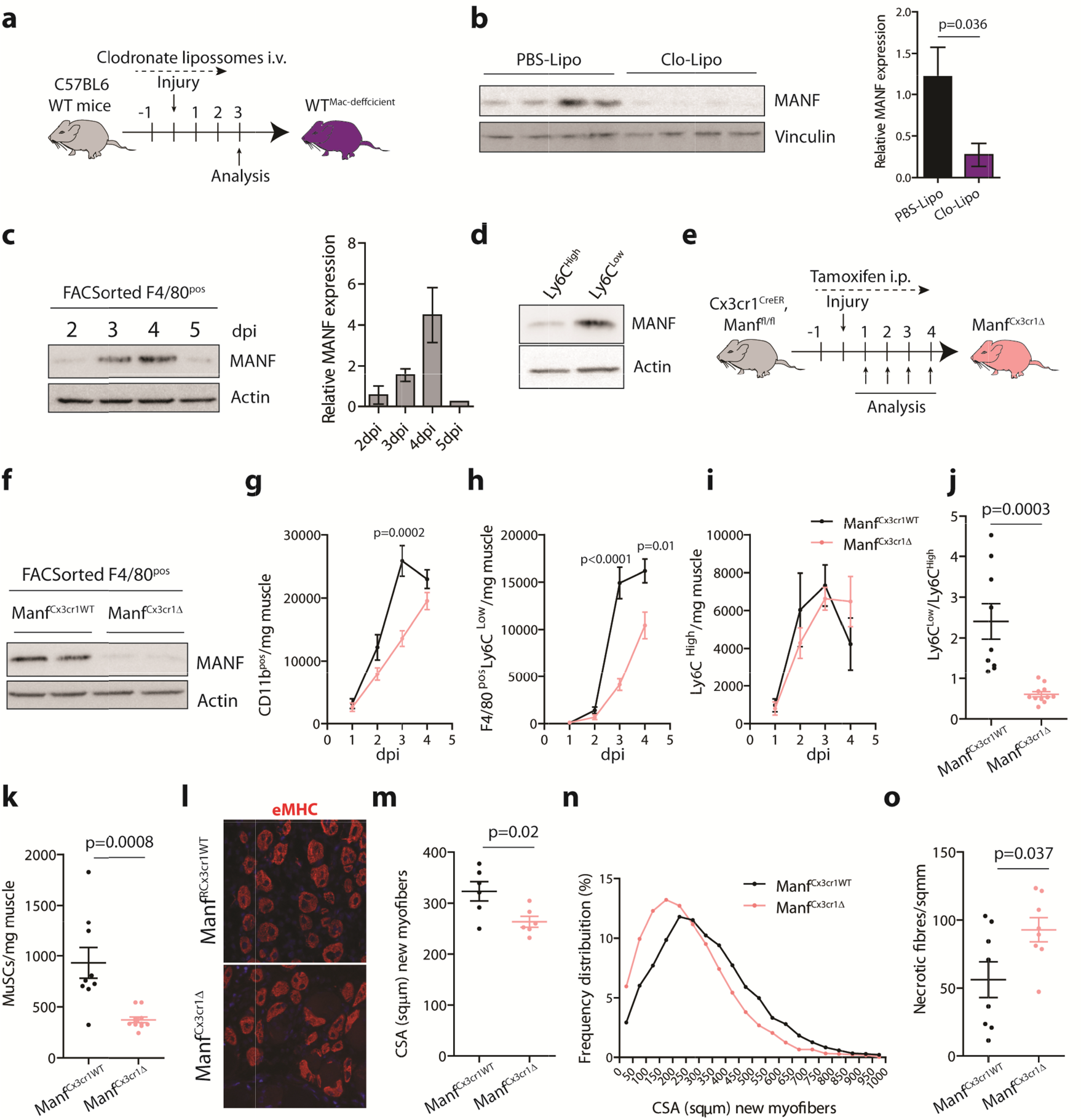
MANF derived from pro-repair macrophages is essential for skeletal muscle regeneration. **a**, Experimental timeline for analysis of animals treated with clodr**o**nate liposomes. **b**, Western blot analysis of MANF levels in protein extracts of QC muscles from wt (C57BL/6) mice, at 3dpi, treated with clodronate liposomes (or control PBS liposomes). Quantification of relative average levels of MANF, normalized to Vinculin, is represented for each condition (n=6/condition). **c**, Western blot analysis of MANF evels in protein extracts from F4/80^pos^ cells FACS-isolated from QC muscles at different time points following injury. Q**u**antification of relative average levels of MANF, normalized to Actin, is represented for each time-point (n=2, 2 and 3dpi; n=3, 4dpi; n=1 (pooled from 4 mice), 5dpi). **d**, Western blot analysis of MANF levels in protein extracts from macrophage subpopulations (Ly6C^High^ and Ly6C^Low^) FACS-isolated from QC muscles of wt mice, at 3dpi. Actin was used to verify equal protein loading in each sample. **e**, Experimental timeline for analysis of animals with conditional ablation of MANF in Cx3cr1-expressing macrophages. **f**, Western blot analysis of MANF levels in protein extracts from F4/80^pos^ cells FACS-isolated from QC muscles of Manf^Cx3cr1WT^ and Manf^Cx3cr1?^ mice at 3dpi. Actin was used to verify equal protein loading in each sample. **g-i**, Quantification, by flow cytometry, of myeloid cells (CD11b^pos^, g), pro-repair macrophages (F4/80^pos^Ly6C^Low^, h) and pro-inflammatory macrophages (Ly6C^High^, i) in regenerating QC muscles of Manf^Cx3cr1WT^ and Manf^Cx3cr1?^ mice at different time points following injury (n=6-11/condition). **j**, Ratio of pro-repair to pro-inflammatory macrophages in regenerating QC muscles of Manf^Cx3cr1WT^ and Manf^Cx3cr1?^ at 3dpi (n=9-11/condition). **k**, Quantification by flow cytometry of MuSCs in regenerating QC muscles of Manf^Cx3cr1WT^ and Manf^Cx3cr1?^ mice at 3dpi (n=9-11/condition). **l**, Representative images of eMHC (red) staining in cryosections of regenerating TA muscle from Manf^Cx3cr1WT^ and Manf^Cx3cr1?^ mice, at 4dpi. DAPI is used to identify nuclei. Quantifications of this staining, for independent animals, are shown in Fig. 2m-n. **m-n**, Quantification of the average cross-sectional area of eMHC+ new myofibres (m) and frequency distribution of new myofibres by size (n), in regenerating TA muscles from Manf^Cx3cr1WT^ and Manf^Cx3cr1?^ mice at 4dpi (n=6/condition). **o**, Quantification of the number of necrotic fibres in regenerating TA muscles from Manf^Cx3cr1WT^ and Manf^Cx3cr1?^ mice at 4dpi (n=8/condition, see Extended data Fig. 2i for representative images). Data are represented as average ± s.e.m. and each n represents one animal. p values are from two-tailed Student’s t-test. dpi, days post-injury; i.v., intravenous injection; PBS-Lipo, PBS Liposomes; Clo-Lipo, Clodronate Liposomes; FACS, Fluorescence activated cell sorting; i.p., intraperitoneal injection; eMHC; embryonic myosin heavy chain; CSA, cross-sectional area; MuSCs, muscle stem cells.

To confirm that this population is the main source of MANF during muscle regeneration we generated a mouse model to selectively deplete MANF in the emerging population of pro-repair macrophages that specifically express Cx3cr1^24^. Indeed, tamoxifen-treated Cx3cr1-Cre^ER^, Manf^fl/fl^ mice (herein referred to as Manf^Cx3cr1Δ^), displayed a complete ablation of MANF protein within the F4/80^pos^ population of macrophages (Fig. 2e-f) and an 80% reduction in MANF levels in whole muscles (Extended data 2e) when compared to oil treated mice (Manf^Cx3cr1WT^). These data support the idea that MANF induction following muscle injury is not only due to an influx of macrophages to the site of injury, but mostly, to a specific increase in MANF protein levels associated with the emergence of pro-repair macrophages within the injured muscle.

### MANF ablation in pro-repair macrophages impairs muscle regeneration

To understand the consequences of MANF loss in the population of pro-repair macrophages we analyzed Manf^Cx3cr1Δ^ animals on a time course following muscle injury, evaluating the repair-associated myeloid response and the efficiency of regeneration. Analysis of myeloid cell populations in Manf^Cx3cr1Δ^ mice revealed alterations in the dynamics of the phenotypic transition between macrophage phenotypes and a reduced presence of myeloid cells within the skeletal muscle (Fig. 2g-j). These alterations were characterized by a significant reduction in the number of pro-repair macrophages (p<0.0001 at 3dpi, Fig. 2h), and no alterations in the numbers of pro-inflammatory macrophages (Fig. 2i), causing a reduction in the ratio of pro-repair to pro-inflammatory macrophages at 3 dpi (Fig. 2j). Interestingly, these defects were not detected at 1dpi and developed primarily between 2dpi and 3dpi, suggesting that they are associated with specific mechanisms operating during the process of phenotypic transition of macrophage phenotypes. Importantly, changes in macrophage phenotypic transition were not due to the tamoxifen treatment, as they were still observed when Manf^Cx3cr1?^ mice were compared with tamoxifen-treated Manf^fl/fl^ mice as controls (Extended data 2f-h).

Evaluation of regenerating muscles at 3 and 4 dpi revealed that MANF ablation in pro-repair macrophages was sufficient to cause defects in muscle repair, evidenced by a reduction in the number of MuSCs present at 3dpi (Fig. 2k), reduced CSA of new myofibres (Fig. 2l-n) and increased presence of necrotic fibres at 4dpi (Fig. 2o and Extended data 2i). Thus, MANF-derived from pro-repair macrophages is essential for a regulated myeloid response and successful debris clearance following muscle injury and effective muscle regeneration.

### MANF is essential for a timely phenotypic transition of macrophages into the pro-repair state

To explore the potential role of MANF in macrophage’s phenotypic transition *ex vivo*, we generated an additional mouse model where MANF is ablated in macrophages in a tamoxifen-independent manner (Manf^LysM?^ mice). Similarly to Manf^Cx3cr1?^ mice, Manf^LysM?^ mice had lower numbers of myeloid cells within the regenerating muscle and a delayed transition between macrophage phenotypes when compared to Manf^fl/fl^ mice (Extended data 3a-d).

To evaluate a specific role of MANF in macrophages’ phenotypic transition, we developed an *ex vivo* assay that allows us to follow the transition of pro-inflammatory macrophages into pro-repair macrophages, independently of the alterations in myeloid cell numbers. In this assay, single cell suspensions isolated from 2dpi muscles were cultured for 16h and the distribution of macrophage populations was quantified by flow cytometry at 0h and 16h (Fig. 3a). Cultured macrophages recapitulated the phenotypic transition observed *in vivo* (Extended data 3e) and Manf^LysM?^ displayed a significant reduction in the emergence of pro-repair macrophages (p<0.05, Fig. 3b and Extended data 3e). Interestingly, these alterations reflected a defect in the early stages of the transition process, as the Ly6C^Int^ population of macrophages was reduced (Extended data 3f) and the transition between Ly6C^Int^ and Ly6C^Low^ was not affected (Extended data 3g). Moreover, in this assay we did not detect significant differences in the number of Cd11b^pos^ cells present in the culture at 16h, when compared with 0h (Extended data 3h), in any of the genotypes, suggesting that the defects cannot be attributed to cell death during the process. Importantly, these defects could be rescued by supplementing recombinant MANF (rMANF) protein in the culture media (Fig. 3c and Extended data 3i), suggesting a feed-forward autocrine mechanism by which MANF derived from pro-repair macrophages sustains the formation of new pro-repair macrophages.

**Figure 3.**
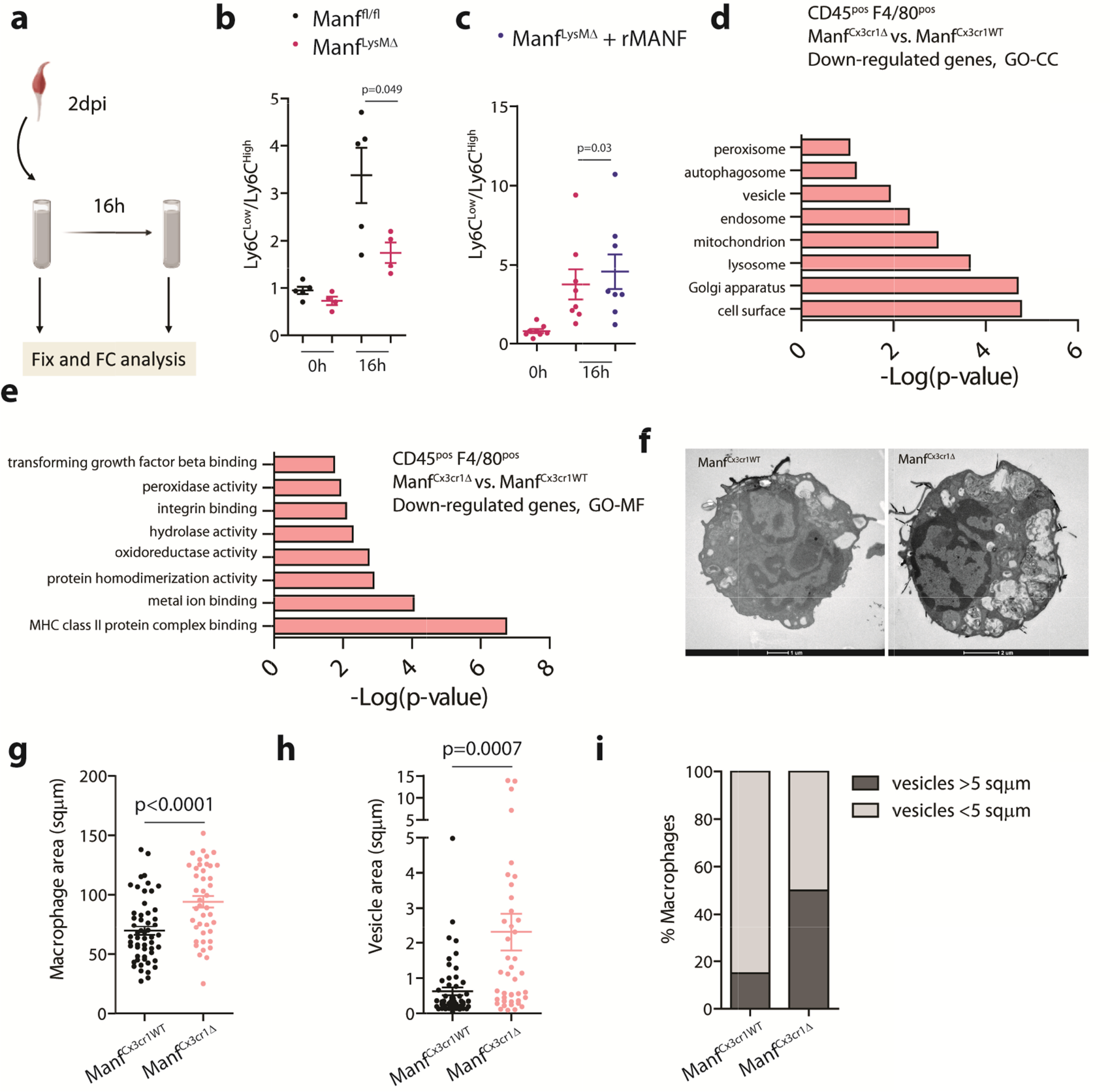
MANF-deficient macrophages have a delayed phenotypic transition and altered structural properties. **a**, Experimental set up for *ex vivo* analysis of single cell suspensions isolated from injured QC muscles. **b**,**c**, Ratio of pro-repair to pro-inflammatory macrophages, quantified by flow cytometry, in single cells suspensions isolated from QC muscles of Manf^fl/fl^ and Manf^LysM?^ mice at 2dpi, 0h and 16h after culture, with or without recombinant MANF supplementation (b, n=4-5/condition; c, n=8/condition). **d**,**e**, GO categories of cellular components (d) and molecular functions (e) showing significant enrichment in the dataset of genes down-regulated in macrophages (CD45^pos^F4/80^pos^) FACS-isolated at 3dpi from QC muscles of Manf^Cx3cr1?^ mice compared to Manf^Cx3cr1WT^ mice (fold change<0.75 and p≤0.05, p values from two-tailed Student’s t-test, n=3/condition). **f**, Representative images of pro-repair macrophages (F4/80^pos^Ly6C^Low^) F CS-isolated at 3dpi from QC muscles of Manf^Cx3cr1WT^ and Manf^Cx3cr1?^ mice, analyzed by TEM. Scale bars: 1μm (Manf^Cx3cr1WT^) and 2μm (Manf^Cx3cr1?^). Quantifications of these images, for independent cells, are shown in Fig. 3g-i. **g-i**, Quantification of macrophage area (g), intracellular vesicle area (h), and percentage of pro-repair macrophages with at least one vesicle bigger than 5 sqμm (i), in a population of pro-repair macrophages FACS-isolated at 3dpi from QC muscles of Manf^Cx3cr1WT^ and Manf^Cx3cr1?^ mice (n=54 macrophages, Manf^Cx3cr1WT^ and n= 42 macrophages, Manf^Cx3cr1?^). Data are represented as average ± s.e.m. and each n represents one animal unless otherwise specified. p values are from two-tailed Student’s t-test. dpi, days post-injury; FC, flow cytometry; rMANF; recombinant Mesencephalic astrocyte-derived neurotrophic factor; GO, Gene Ontology; CC, cellular component; MF, molecular function;

RNA sequencing (RNAseq) analysis of macrophages (CD45^pos^F4/80^pos^) freshly isolated from Manf^Cx3cr1?^ and Manf^Cx3cr1WT^ muscles at 3dpi revealed changes in key cellular processes (Fig. 3d-e and Extended data 4a). Gene ontology analysis of the dataset of down-regulated genes revealed enrichment for genes associated with lysosomal and endosomal compartments (Fig. 3d), and with molecular functions related to hydrolase, peroxidase and oxidoreductase activity (Fig. 3e). Consistently, functional classification of all differentially expressed genes revealed alterations in the biological processes of MHC class II antigen presentation, a cellular process dependent on the efficient lysosomal digestion of internalized proteins by phagocytosis, and in the response to oxidative stress (Extended data 4a). Additionally, functional classes of biological functions associated with normal functions of pro-repair macrophages were also altered in Manf-deficient macrophages (Extended data 4a), suggesting functional alterations in the context of tissue repair.

To better characterize the alterations of Manf-deficient macrophages we zoomed in on the pro-repair population. The F4/80^pos^Ly6C^Low^ subpopulation of pro-repair macrophages was isolated by FACS from Manf^Cx3cr1?^ and Manf^Cx3cr1WT^ muscles at 3dpi and analyzed by transmission electron microscopy (TEM). Manf-deficient pro-repair macrophages exhibited marked structural differences (Fig. 3f), characterized by a significant increase in size (p<0.0001, Fig. 3f-g) and an accumulation of large vesicular structures, often filled with undigested cellular material (Fig. 3f and 3h-i). These structural alterations are consistent with the gene expression changes associated with alterations in the lysosomal compartment and hydrolytic activity and may reflect alterations in the phagocytic pathway, previously associated with the phenotypic transition of macrophages into a pro-repair state^13^.

### MANF restores the repair associated myeloid response and improves muscle regeneration in ageing

To understand if similar defects were present in old animals, we performed RNAseq analysis of pro-repair macrophages (CD11b+F4/80^pos^Ly6C^Low^) isolated from aged muscles at 3dpi. The dataset of up-regulated genes was enriched for gene ontologies associated with inflammatory activation, suggesting a shift in the gene expression profile of the pro-repair population towards a pro-inflammatory phenotype (Fig. 4a). Similarly to what we observed in MANF-deficient macrophages, the dataset of down-regulated genes revealed enrichment in gene ontologies of cellular components associated with lysosomal and endosomal compartments, along with changes associated with filopodia and lamellipodia (Fig. 4b). Genes down-regulated in aged macrophages also classified in biological processes with relevance within the context of tissue regeneration, such as wound healing, endothelial cell morphogenesis or regulation of cell-cell adhesion (Extended data 4b).

**Figure 4.**
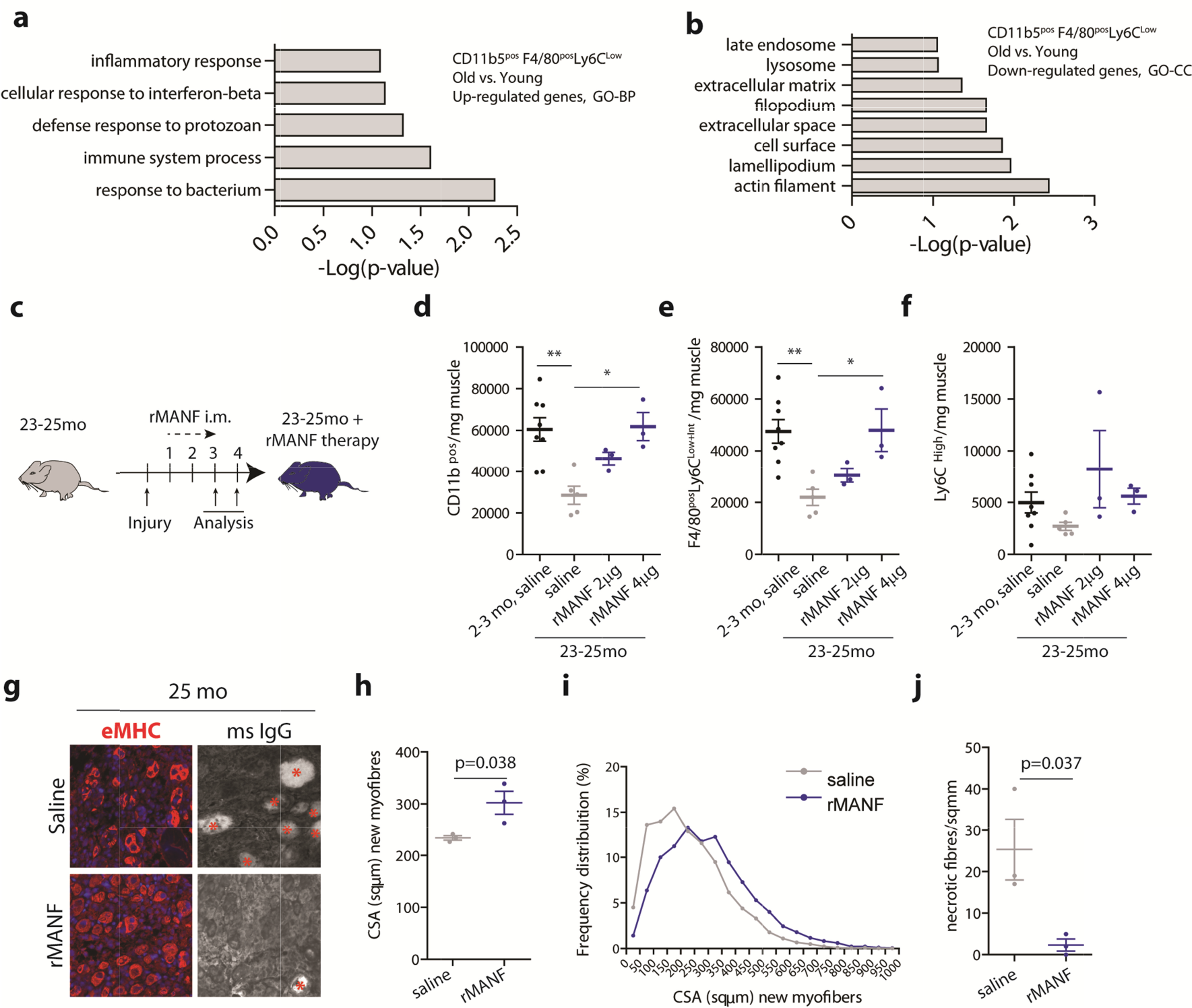
MANF therapy restores the repair-associated myeloid response and regenerative success in aged skeletal muscles. **a**,**b**, GO categories of biological processes (a) and cellular components (b) showing significant enrichment in the dataset of genes up-regulated (**a**) or down-regulated (b) in pro-repair macrophages (F4/80^pos^Ly6C^Low^) FACS-isolated at 3dpi from QC m**u**scles of old (22-24mo) mice compared to yg (2-6mo) mice (fold change<0.75 or >1.5 and p≤0.05, p values from two-tailed Student’s t-test, n=3/condition). **c**, Experimental timeline for analysis of aged animals after receiving intramusc lar injections of rMANF protein. **d-f**, Quantifi**c**ation, by flow cytometry, of myeloid cells (CD11b^pos^, d), pro-repair macrophages (F4/80^pos^Ly6C^Low+Int^, e) and pro-inflammatory macrophages (Ly6C^High^, f) in regenerating TA muscles of yg (2-3mo) and old (23-25mo) wt (C57BL/6) mice at 3dpi, treated with i.m. injections of saline or Rmanf (n=3-8/condition). **g**, Representative images of eMHC (red, left) or mouse IgG (right) staining in cryosections of regenerating TA muscles from old (25mo) mice, at 4dpi, treated with intramuscular injections of rMANF or saline. DAPI (blue) is used to identify nuclei. Asterisks indicate necrotic myofibres. Quantifications of these stainings, for independent animals, are shown in Fig. 4h-j. **h-j**, Quantification of the average cross-sectional area of new myofibers (h), frequency distribution of new myofibres by size (i) and average number of necrotic fibres (j) in regenerating TA muscles from old (25mo) mice, at 4dpi, treated with intramuscular injections of rMANF or saline (n=3/condition). Data are represented as average ± s.e.m. and each n represents one animal. In d-f, p values are from one-way ANOVA with Bonferroni’s multiple comparison post-test. In h, j, p values are from two-tailed Student’s t-test. GO, Gene Ontology; BP, biological process; CC, cellular component; i.m., intramuscular injection; rMANF; recombinant Mesencephalic astrocyte-derived neurotrophic factor; eMHC; embryonic myosin heavy chain; ms IgG, mouse Immunoglobulin; CSA, cross-sectional area.

Since MANF induction following injury was impaired in aged animals (Fig. 1c and Extended data Fig. 1b-c); defects in the repair-associated myeloid response and muscle regeneration were similar in aged and MANF-deficient mice (Fig. 1d-r); and MANF supplementation was sufficient to restore pro-repair macrophages in models of MANF deficiency (Fig. 3c), we sought to explore whether MANF therapy could allay the age-related defects in muscle regeneration. Our strategy consisted in delivering rMANF through daily intramuscular (i.m.) injections (2 or 4ug/ injections) starting at 1 dpi and up to the day of analysis (Fig. 4c). We found that delivery of rMANF to injured muscles of aged mice was sufficient to normalize the repair-associated myeloid response, restoring the numbers of myeloid cells and pro-repair macrophages at 3 dpi (Fig. 4d-f). The effects were dose-dependent and MANF therapy at 4ug/i.m. injection resulted in a complete rescue of the repair-associated myeloid response. Importantly, the same regimen of MANF therapy improved muscle regeneration in aged animals, resulting in an increase in the CSA of new myofibres and a reduction in the accumulation of uncleared necrotic fibres (Fig. 4g-j). These data show that restoring MANF levels during regeneration in aged muscles is sufficient to normalize the repair-associated myeloid response and suggest that immune modulatory interventions can be used to improve the regenerative capacity of aged muscles.

## Discussion

Age-related alterations in myeloid populations occur in the aged skeletal muscle, in homeostasis, with detrimental consequences for MuSC activity and tissue health^25-28^. However, the effects of ageing on the myeloid response associated with muscle regeneration have just started to be explored^28,29^. Here we uncovered a new regulator of the myeloid response to muscle injury, MANF, involved in the maintenance of a proper balance of macrophage populations throughout muscle regeneration. By identifying a critical down-regulation of MANF signalling in the aged regenerating skeletal muscle, we demonstrated, for the first time, how age-related changes in immune modulatory mechanisms contribute to the skeletal muscle’s regenerative failure in ageing.

The MANF-deficient models used in this study recapitulate several defects of muscle regeneration present in aged animals, including the decreased capacity to clear necrotic muscle fibres. Studies in recent years have described multiple mechanisms regulating macrophage phenotypic transition during muscle regeneration^12^. Age-related impairments of the myeloid system have been identified in other contexts, including metabolic and phagocytic defects^30,31^. Our analysis points to defects in the intracellular processing of cellular debris phagocytized by macrophages in conditions of MANF deficiency, suggesting a link between altered macrophage phagocytic function and dysregulation of macrophage responses in ageing. Considering the increased severity of the regenerative impairments in the full MANF knock out, relative to models of macrophage-specific ablation, and the ability of delivered rMANF to rescue functional defects in MANF-deficient and aged animals, it is likely that MANF extracellular activity is important in this regulatory process.

The use of immune modulatory adjuvants in regenerative medicine can be an attractive strategy to improve the clinical outcome of stem-cell based therapies in the elderly^1^. The implementation of these therapeutic solutions is going to require a better understanding of the age-related impairments of immune mechanisms associated with tissue repair. From this work, MANF emerges as a potential candidate to be used in a clinical setting where MuSCs are applied to the aged skeletal muscle.

## Acknowlegements

We would like to thank the Flow Cytometry, Comparative Pathology, Bioimaging and Rodent facilities of Instituto de Medicina Molecular João Lobo Antunes for their technical support. We thank A.S. Pacheco and E.M. Tranfield from the Electron Microscopy Facility at the Instituto Gulbenkian de Ciência for sample processing, data collection, and discussions about the results. This work was suported by EMBO (IG4448 to PSV) and FCT (PTDC/MED-OUT/8010/2020 and EXPL/MED-OUT/1601/2021 to PSV and JN). PSV was supported by “la caixa” Foundation Junior Leader Fellowship (LCF/BQ/PI19/11690006). JN was supported by an assistant research contract from FCT (2021.03843.CEECIND).

## Authors Contribuitions

JN and PSV conceived the study, designed experiments, analyzed and interpreted data and wrote the manuscript. NSS and MFB designed and performed experiments, analyzed and interpreted data. IBA performed experiments and analyzed data. PL performed the ELISA analysis of muscle extracts. All authors revised the manuscript.

## Data Availability

RNAseq datasets will be made available upon publication.

## Methods

### Animals

All mice used in these studies were housed at the DGAV accredited rodent facility of Instituto de Medicina Molecular, in individually ventilated cages within a specific and opportunistic pathogen-free (SPOF) facility, on a standard 12/12h light cycle. The care and use of experimental animals complied with relevant institutional and national animal welfare laws, guidelines and policies.

Old wt C57BL/6 mice were purchased from Charles River, Europe with 18-20 months and further aged in house until analysis. Young wt mice were either purchased from Charles River, Europe; or born in house and generated using C57BL/6 breeders purchased from Charles River, Europe.

To create a mouse model allowing an inducible and ubiquitous ablation of *Manf* we generated Rosa26^CRE-ERT/+^Manf^fl/fl^ mice. Mice carrying the Rosa26^CRE-ERT^ allele are B6;129-Gt(ROSA)26Sortm1(cre/ERT)Nat/J and were purchased from The Jackson Laboratory (JAX, stock no: 004847). In these mice a Cre^ERT^ cassette is inserted within intron 1 of the GT(ROSA)26Sor locus and is expressed under the control of its endogenous promoter. Mice carrying the Manf^fl^ allele were previously described^20^. Heterozygous carriers of the Cre^ERT^ allele and homozygous carriers of the Manf^fl^ allele were used for these studies. Cx3cr1^CRE-ER/+^, Manf^fl/fl^ mice were previously described^20^. To induce Cre activity, mice received daily intra-peritoneal injections of tamoxifen (T5648-1G – Sigma) in sterile corn oil, at a dose of 75mg/Kg of body weight. Control mice were littermates of the same genotype that received sham injections (corn oil) or Manf^fl/fl^ mice that received tamoxifen injections. To create a mouse model where MANF is ablated in macrophages in a tamoxifen-independent manner we generated LysM^CRE/+^/MANF^fl/fl^ mice. Mice carrying the LysM^CRE^ allele are B6.129P2-Lyz2tm1(cre)Ifo/J and were purchased from JAX (stock number 004781). Heterozygous carriers of the Cre allele and homozygous carriers of the Manf^fl^ allele were used for these studies. Control mice were MANF^fl/fl^ littermates without the Cre allele. LysM^CRE/+^, Manf^fl/fl^; Cx3cr1^CRE-ER/+^, Manf^fl/fl^ mice; and MANF^fl/fl^ mice were generated at the Buck Institute for Research on Aging (Novato, CA, USA) and re-derived into SPOF C57BL/6J strain by *in vitro* fertilization. Primers used for genotyping all mouse strains are listed in Supplementary table 1.

### In vivo procedures in mice

All procedures involving animals were approved by Direção Geral da Alimentação e Veterinaria (DGAV) and performed at the rodent facility of Instituto de Medicina Molecular.

### Induction of muscle regeneration

Regeneration of skeletal muscle was induced by intramuscular (i.m.) injection of sterile 1.2% Barium Chloride (Sigma: 342920) in saline solution (0.9% NaCl, B. BRAUN) into the tibialis anterior (TA, 40ul) or quadriceps (QC, 50ul) muscle of the mice. At the designated time points after injury, mice were euthanized and muscles were collected for analysis.#Animals were anesthetized for the procedure with Isoflurane inhalation. Tamoxifen injected mice were anesthetized with Ketamine 75mg/Kg of body weight + Medetomidine 1mg/Kg of body weight before i.m.injection.

### Macrophage ablation

Chemical ablation of macrophages was performed using a clodronate-liposome solution. Clodronate-liposomes or PBS-liposomes (LIPOSOMA – Research solution SKU: CP-010-010) at 5mg/ml were inject intravenously (tail) at a dose of 100ul/10g of body weight. Animals received one injection on the day before injury, and then daily injections until analysis starting at 1dpi.

### rMANF intramuscular injection

Injured mice received daily i.m. injections of 20ul of saline solution (0.9% NaCl, B. BRAUN) containing 2μg or 4μg of hrMANF protein (P-101-100, Icosagene), in the injured muscle starting at 1 dpi until the day of analysis. Control animals in the same experiment received the same regimen of i.m. injections of saline solution without rMANF supplementation.

### Histological analysis, imaging and quantification methods for muscle tissue

#### Muscle tissue harvesting and storage

Animals were euthanized and tissues were harvested. For histology analysis the dissected TA muscle was mounted vertically on a tragacanth gum (G1128-100G – Sigma) placed on a card board. The tissue was frozen in methyl-butane (VWRC103614T – VWR) cooled with liquid nitrogen, for about 13 to 17 seconds depending on muscle size and stored at -80ºC for further analysis as described^32^. Frozen tissue was cryosectioned at 10μm thickness on a cryostat (LEICA CM 3050S), collected on Superfrost microslides (VWR) and stored at -80ºC until analysis. Samples processed for RNA and protein analysis were flash frozen in cryotubes submerged in liquid nitrogen.

#### H&E staining of muscle sections

#Muscle cryosections collected on slides were thawed at room temperature (RT) for 10min. Cryosections were placed in distilled water for 5min, stained on Harris Hematoxylin (05-06004E – Enzifarma), placed under running water for 5min, dipped on ethanol 70%, stained on eosin (HT110132-1L-Sigma). Stained tissue was serially dehydrated in 70%, 95% and 100% (twice) ethanol for 30 sec on each alcohol, incubated in xylene (3803665EGDG – Leica Microsystems) for at least 10 min and mounted with MICROMOUNT (3801731DG - Leica Microsystems).

#### Immunohistochemistry (IHC) of muscle cryosections and nuclei staining

Muscle cryosections collected on slides were thawed, permeabilized with PFA 4% in Phosphate-buffered saline (PBS) 10 min at RT, incubated in boiling 10mM Citrate buffer 45 min, blocked with Mouse on Mouse Blocking Reagent (R&D Systems) 2h and incubated with primary antibody, diluted in blocking solution, overnight (O/N) at 4°C. Primary antibody was washed 4x with PBS containing 0.1% Tween20 (PBS-T) and detected by incubating 2h30min with Alexa conjugated secondary antibodies (Abcam). Secondary antibody was washed 5x with PBS-T. Nuclei were stained for 5min with 300nM DAPI (4’,6-diamidino-2-phenylindole) in PBS at RT. Slides were rinsed in PBS and mounted with Mowiol mounting media and microscope cover glass No. 1.5H (Marienfeld). Staining of necrotic myofibres using secondary antibody anti-mouse IgG coupled to Alexa-647 was performed without primary antibody incubation overnight. Co-staining of F4/80 and MANF was performed without the permeabilization with Citrate buffer and blocked with Horse Serum (HS) 10% in PBS-T. For information on primary antibodies used see Supplementary Table 2.

#### Imaging

Digital images were acquired at the Bioimaging and Comparative Pathology facilites of Instituto de Medicina Molecular using: (1) a digital slide scanner NanoZoomer SQ (HAMAMATSU), with an objective of 20X magnification, for H&E stained sections; (2) a motorized inverted widefield fluorescence microscope (Zeiss) equipped with CCD camera (Photometrics CoolSNAP HQ CCD), with a 20X objective for IHC stained sections (eMHC, IgG). The total stained sections after imaging were reconstructed through image overlay (10%); (3) a Zeiss LSM 710 confocal laser scanning microscope (F4/80/MANF IHC).

#### Image quantification

To assess the effectiveness of the muscle regeneration the individual new myofibres (eMHC^pos^) were manually outlined in the total muscle section and their cross-sectional area (CSA) was determined with the public domain image analysis software ImageJ. The number of necrotic fibres in total muscle section was quantified using the same software.

#### Flow cytometry (FC) analysis

To obtain single cell suspensions, muscles were mechanically disaggregated and dissociated in DMEM 1% P/S media containing collagenase B (Roche) 0.2% and Calcium dichloride (CaCl_2_) 0.5 mM at 37 °C for 1h and then filtered through 70 μm cell strainers (Falcon). Cells were incubated in 1x Red Blood Cell (RBC) lysis buffer (Santa Cruz Biotechnology) for 10 min on ice, resuspended in 1ml of DMEM 10% FBS 1% P/S media and counted. For flow cytometry analysis (FC analysis), single cell suspension samples were resuspended in PBS containing 5% HS with fluorophore-conjugated antibodies at a density of 1×10^6^ cells/100 μl, incubating 30 min at 4ºC, protected from light. Cells were re-suspended in PBS containing 5% HS for FC analysis. For information on antibodies used see Supplementary Table 3. CD45, CD31 markers were used to exclude the Lin (-) negative population from single live cell population and the population of MuSCs and FAPS were identified as α7-integrin^pos^ and Sca-1^pos^, respectively. Live cells were identified using LIVE/DEAD™ FixableNear-IR Dead Cell Stain (Invitrogen). Gating strategy used in FC analysis of CD45^pos^ immune cell population, endothelial cells, FAPS and MuSCs is shown in Extended data Figure 1f. Gating strategy used in FC analysis of myeloid cells (CD11b^pos^), pro-repair macrophages (F4/80^pos^Ly6C^Low^), pro-inflammatory macrophages (Ly6C^High^), and neutrophils (Ly6G^pos^) is presented in Extended data Figure 1i. Characterization of cell populations was performed at the Flow cytometry facility of Instituto de Medicina Molecular, using a cell analyzer LSRFortessa X-20 (BD Bioscience) with FACSDiva 8.0 software. Flow cytometry data were analyzed using FlowJo (BD Biosciences) analysis software.

#### Fluorescence activated cell sorting (FACS) of macrophage populations

Single cell suspensions, obtained as described above, were used to isolate macrophage populations through staining with fluorochrome-conjugated antibodies presented in Supplementary Table 3. CD45^pos^ F4/80^pos^ macrophages were selected from the viable cells present in the single cell suspension. F4/80^pos^Ly6C^Low^ and LyC6^High^ macrophages were isolated using the gating strategy presented in Extended data Figure 1i. The isolation of pure populations of cells was performed at the Flow cytometry facility of Instituto de Medicina Molecular using a FACSAria IIu (BD Bioscience) or a FACSAria III (BD Bioscience) using the software FACSDiva 6.1.3. Cells were collected in PBS containing 5% HS and used either for protein extraction, RNA extraction or TEM analysis.

#### Ex-vivo macrophage analysis

Single cell suspensions of 2pi injured muscles were obtained as described above. For each animal, 500 000 cells were collected at 0h or cultured and collected after 16h. Cells were incubated in suspension at 37ºC in SF medium (Corning® SF Medium, with L-glutamine and 1 g/L BSA) supplemented with 10% FBS and 1% Pen/Strep. In conditions of MANF supplementation, rMANF (P-101-100, Icosagene) was used at a concentration of 10μg/ml. Cells collected at 0h and 16h were stained for FC analysis of muscle immune populations as described above.

### Transmission Electron Microscopy analysis of macrophages

#### Sample processing and imaging

Pro-repair macrophages were isolated by FACS as described above, plated on 12 mm coverslips inserted in a 24-well plate with DMEM media containing 1% PS and 10%FBS, and allowed to adhere for 2h. Sample processing and imaging was performed at the Electron Microscopy facility at Instituto Gulbenkian de Ciênca (IGC, Lisbon, Portugal). Cells were fixed with 2% formaldehyde (FA) - 2.5% Glutaraldehyde in 0.1M phosphate buffer (PB) for 45min on ice and then fixed O/N in 1% FA in PB at 4°C. The following day, cells were washed 2x in PB, post-fixed with 1% osmium in PB for 1h on ice, washed 2x in PB, 2x in water, stained with 1% tannic acid 20min on ice, washed 2x in water, stained with 0.5% in Uranyl acetate 1h at RT and serial dehydrated in increasing concentrations of ethanol. Coverslips with cells were mounted on top of EPON capsules and baked at 60°C O/N. Sections of 70 nm were obtained using a UC7 Ultramicrotome (Leica) and stained with uranyl acetate and lead citrate for 5 minutes each. Images of single macrophages were acquired on a Tecnai G2 Spirit BioTWIN Transmission Electron Microscope (TEM) from FEI operating at 120 keV and equipped with an Olympus-SIS Veleta CCD Camera.

#### Image quantifications

Quantification of individual pro-repair macrophages from Manf^Cx3cr1Δ^ (n=42) and Manf^Cx3cr1WT^ (n=54) mice, analyzed by TEM, was performed using the ImageJ software, after scale normalization. Intracellular vesicles were manually surrounded, and the number and area were determined. In addition, the area of each cell was also calculated.

### Protein analysis

#### Preparation of muscle protein extracts

Whole muscle samples, or cells obtained from FACS, were homogenized in Lysis buffer (50 mM Tris pH 7.5, 150 mM NaCl, 0.5% NP-40, 5 mM EDTA, 1%Triton, in Mili-Q water) supplemented with protease inhibitors and phosphatase inhibitors (Sigma) for 45 min or 20 min at 4ºC, respectively. The supernatant protein extracts were recovered by centrifugation and protein concentration in samples was determined using Bradford Reagent (VWR).

#### Western blot

Western blot analyses were performed on 12% SDS-PAGE. After electrophoretic separation, proteins were transferred onto nitrocellulose membranes using a Trans Blot Turbo Transfer system (BioRad). Membranes were blocked with Tris-buffered saline-0.1% Tween 20 (TBS-T) containing 5% milk for 1h and incubated overnight at 4ºC with primary antibodies. Membranes were then incubated 1h with a peroxidase-conjugated secondary antibody (1:10000; Abcam), and developed using Pierce ECL Western blotting substrate (ThermoScientific) or Clarity™ Western ECL Substrate (BioRad). For information on primary antibodies used see Supplementary Table 4. Ponceau S solution (Sigma) was applied to the nitrocellulose membranes before the blocking step to assess the total protein present.

#### Enzyme-Linked Immunosorbent Assay (ELISA)

MANF concentrations in muscle tissue samples were quantified using an in-lab mouse MANF (mMANF) ELISA^33^. mMANF ELISA recognized both mouse and human MANF but did not recognize MANF homolog CDNF or give signal from tissue lysates from Manf-/-mice, indicating that it was specific for MANF. Dynamic range of mMANF ELISA was 62.5 – 1000 pg/ml and its sensitivity 29 pg/ml. For mMANF ELISA measurement, muscle lysates from young mice were diluted at 1:500 in blocking buffer (1% casein in PBS-0.05% Tween-20), and lysates from young and old mice (2dpi, 3dpi) at 1:200, respectively. Recombinant human MANF (P-101–100; Icosagen) was used as a standard. All samples were measured in duplicate.

### RNA analysis

#### Preparation of RNA samples

Total RNA from frozen muscle samples was extracted using TRIzol (Invitrogen), according to the supplier’s instructions. Total RNA from sorted cells was extracted using RNeasy Micro kit (Qiagen), according to the supplier’s instructions.

#### Reverse Transcription and real-time qPCR (RT-qPCR)

Complementary DNA (cDNA) was synthesized using iScript cDNA synthesis kit (BioRad). Real-time PCR was performed on ViiA 7 Real-Time PCR System (Thermofisher Scientific), using Powerup SYBR Green MASTER MIX (Applied Biosystems). Expression of specific genes in each sample was normalized to beta-actin and results are shown as gene expression levels relative to levels in control samples which are arbitrarily set to one. For information on primer sequences see Supplementary Table 5.

#### RNA sequencing and bioinformatics analysis

Library preparation, RNA sequencing, read mapping and FPKM quantification was performed as a service at Novogene (Cambrige, UK). RNA samples prepared as described above, from sorted macrophages (Manf^Cx3cr1Δ^ vs Manf^Cx3cr1WT^) or pro-repair macrophages (Yg vs. old) were shipped to Novogene. Novogene team was responsible for Illumina library preparation (poly A enrichment) and sequencing using a NovaSeq instrument to generate 150 bp pair-end reads with an output of 6G per sample. Gene Ontology and KEGG analysis was carried out using The Database for Annotation, Visualization and Integrated Discovery (DAVID, https://david.ncifcrf.gov/, ref. 76).

#### Statistical analysis

All data are presented as average and standard error of the mean (s.e.m.). Statistical analysis was carried out using GraphPad Prism 5. For comparisons between two groups, a two-tailed Student’st test was used to determine statistical significance, assuming normal distribution and equal variance. For multiple comparisons, one-way ANOVA with Bonferroni’s multiple comparison post-test were used to determine statistical significance.

