## Extended data and Supplementary material for "Ageing disrupts MANF-mediated immune modulation during skeletal muscle regeneration"

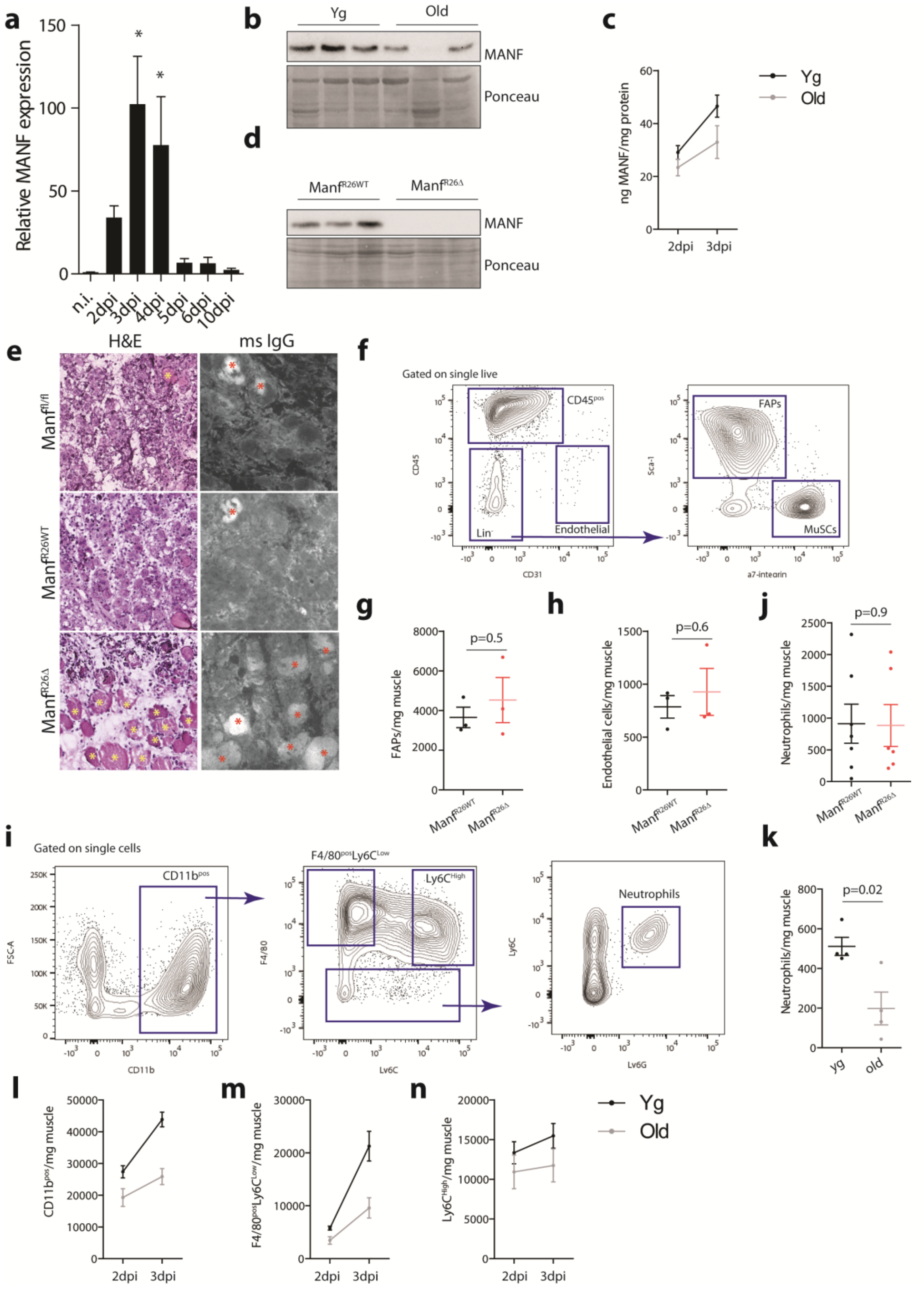
a
Relative MANF expression
150
100
50
0
*
*
n.i.
2dpi
3dpi
4dpi
5dpi
6dpi
10dpi
b
Yg
Old
MANF
Ponceau
c
ng MANF/mg protein
60
40
20
0
2dpi
3dpi
Yg
Old
d
Manf^R26WT
Manf^R26Δ
MANF
Ponceau
e
H&E
ms IgG
Manf^fl/fl
Manf^R26WT
Manf^R26Δ
f
Gated on single live
CD45^pos
Lin⁻
Endothelial
CD45
CD31
FAPs
MuSCs
Sca-1
a7-integrin
g
FAPs/mg muscle
8000
6000
4000
2000
0
p=0.5
Manf^R26WT
Manf^R26Δ
h
Endothelial cells/mg muscle
1500
1000
500
0
p=0.6
Manf^R26WT
Manf^R26Δ
j
Neutrophils/mg muscle
2500
2000
1500
1000
500
0
p=0.9
Manf^R26WT
Manf^R26Δ
i
Gated on single cells
CD11b^pos
FSC-A
CD11b
F4/80^pos Ly6C^Low
Ly6C^High
F4/80
Ly6C
Neutrophils
Ly6G
k
Neutrophils/mg muscle
800
600
400
200
0
p=0.02
yg
old
l
CD11b^pos/mg muscle
50000
40000
30000
20000
10000
0
2dpi
3dpi
m
F4/80^pos Ly6C^Low/mg muscle
30000
20000
10000
0
2dpi
3dpi
n
Ly6C^High/mg muscle
20000
15000
10000
5000
0
2dpi
3dpi
Yg
Old

**Extended data Figure 1. Regulation of muscle regeneration by MANF**

**a,** Relative levels of MANF mRNA, detected by RT-qPCR, in TA muscles of yg wt (C57BL/6) mice non-injured and at different time points following injury (2, 3, 4, 5, 6, 10 dpi) (n=3-4/ time point). **b,** Illustrative Western blot analysis of MANF levels in protein extracts of TA muscles from yg (2-6mo) and old (22-25mo) wt (C57BL/6) mice at 3dpi. Ponceau S-staining of the membrane was used to verify equal protein loading in each sample. **c,** MANF protein levels, quantified by ELISA, in extracts of TA muscles from yg (2-6mo) and old (22-25mo) wt (C57BL/6) mice, at 2 and 3dpi (n=3-5/ condition). **d,** Western blot analysis of MANF levels in protein extracts from $Manf^{R26WT}$ and $Manf^{R26\Delta}$ mice at 3dpi. Ponceau S-staining of the membrane was used to verify equal protein loading in each sample. **e,** Representative images of cryosections from TA muscles of $Manf^{fl/fl}$ , $Manf^{R26WT}$ and $Manf^{R26\Delta}$ mice, at 4 dpi, stained with H&E (left) and immunostained with mouse IgG (right). Asterisks indicate necrotic myofibers. **f,i,** Gating strategy used in flow cytometry analysis of (f) $CD45^{pos}$ immune cell population, endothelial cells, FAPs and MuSCs; and (i) myeloid cells ($CD11b^{pos}$), pro-repair macrophages ($F4/80^{pos}Ly6C^{Low}$), pro-inflammatory macrophages ($LyC6^{High}$), and neutrophils ($F4/80^{neg}Ly6G^{pos}$). **g,h,j,k,** Quantification, by flow cytometry, of FAPS (g, n=3/condition), endothelial cells (h, n=3/condition) and neutrophils (j, n=6-7/condition) in regenerating muscles of $Manf^{R26WT}$ and $Manf^{R26\Delta}$ mice at 3dpi; and neutrophils (k, n=4/condition) in regenerating muscles of yg (2-6mo) and old (22-24mo) wt (C57BL/6) mice at 3dpi. **l-n,** Quantification, by flow cytometry, of myeloid cells ($CD11b^{pos}$, l), pro-repair macrophages ($F4/80^{pos}Ly6C^{Low}$, m) and pro-inflammatory macrophages ($Ly6C^{High}$, n) in regenerating QC muscles of yg (2-6mo) and old (22-25mo) wt (C57BL/6) mice at 2 and 3dpi (n=4-6/condition). Data are represented as average ± s.e.m. and each n represents one animal. In a, p values are from one-way ANOVA with Bonferroni's multiple comparison post-test. In all other graphs, p values are from two-tailed Student's t-test. n.i., non-injured; dpi, days post-injury; H&E, Hematoxylin and Eosin; msIgG, mouse Immunoglobulin; FAPs, Fibroadipogenic progenitors; MuSCs; Muscle Stem Cells; yg, young

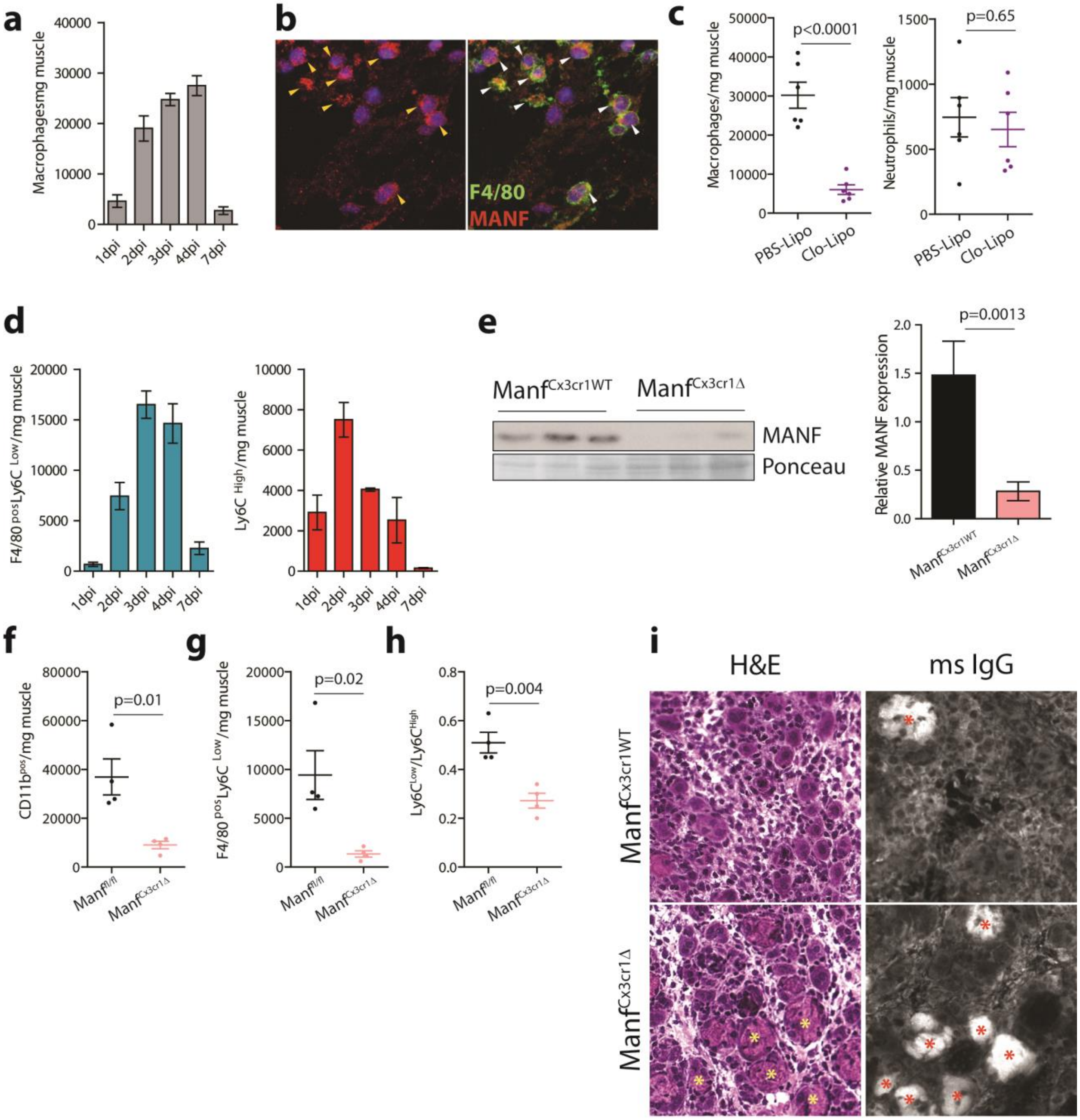


**Extended data Figure 2. Macrophage-derived MANF in muscle regeneration**

**a,d** Quantification, by flow cytometry, of macrophages (a), pro-repair macrophages (d, left; F4/80$^{pos}$Ly6C$^{Low}$) and pro-inflammatory macrophages (d, right; Ly6C$^{High}$) in regenerating TA muscles of wt (C57BL/6) mice at different time points following injury (n=3-8/condition). **b,** Representative images of cryosections from TA muscles immunostained against F4/80 (green) and MANF (red). DAPI is used to identify nuclei. Arrowheads indicate cells with high MANF expression co-localized with F4/80. **c,** Quantification, by flow cytometry, of macrophages and neutrophils in regenerating QC muscles of wt (C57BL6/J) mice at 3dpi, treated with clodronate liposomes or control PBS liposomes (n=6/condition). **e,** Western blot analysis of MANF levels in protein extracts from muscles of $Manf^{Cx3cr1WT}$ and $Manf^{Cx3cr1\Delta}$ mice at 3dpi. Quantification of average relative levels of MANF, normalized to Ponceau S-staining, are represented for each condition (n=5-9/condition). **f-h,** Quantification, by flow cytometry, of myeloid cells ($CD11b^{pos}$, f), pro-repair macrophages ($F4/80^{pos}Ly6C^{Low}$, g), and ratio of pro-repair to pro-inflammatory macrophages ($LyC6^{Low}/LyC6^{High}$, h) in regenerating QC muscles of tamoxifen treated $Manf^{fl/fl}$ and $Manf^{Cx3cr1\Delta}$ mice at 3dpi (n=4/ condition). **i,** Representative images of cryosections from TA muscles of $Manf^{Cx3cr1WT}$ and $Manf^{Cx3cr1\Delta}$ mice, at 4 dpi, stained with H&E (left) and immunostained with mouse IgG (right). Asterisks indicate necrotic myofibers. Data are represented as average ± s.e.m. and each n represents one animal. p values are from two-tailed Student's t-test. dpi, days post-injury; PBS-Lipo, PBS Liposomes; Clo-Lipo, Clodronate Liposomes; H&E, Hematoxylin and Eosin; msIgG, mouse Immunoglobulin.

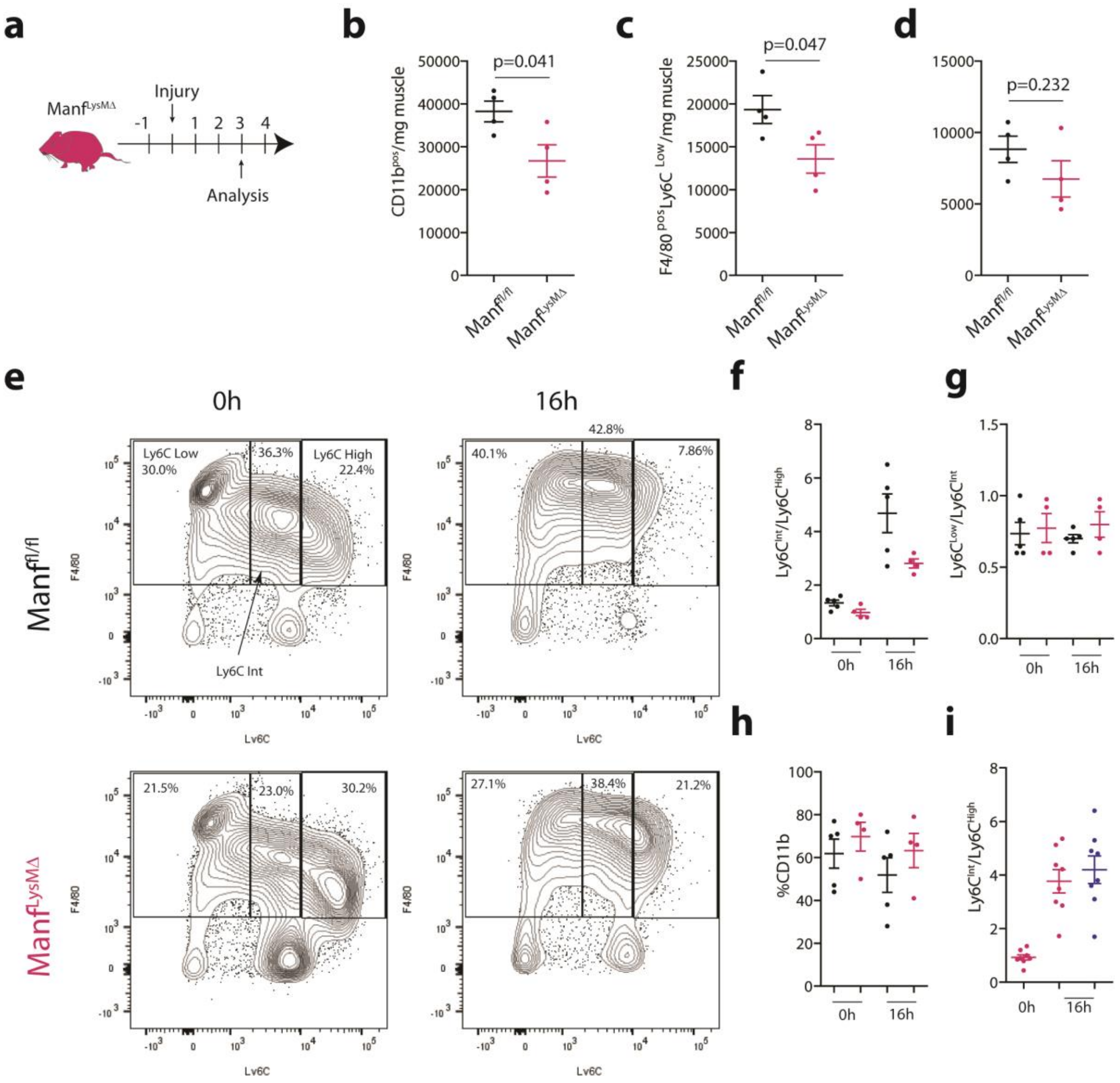


**Extended data Figure 3. MANF-deficiency affects macrophage phenotypic transition**

**a,** Experimental timeline for analysis of animals with ablation of MANF in macrophages. **b-d,** Quantification, by flow cytometry, of myeloid cells (CD11b$^{pos}$, b), pro-repair macrophages (F4/80$^{pos}$Ly6C$^{Low}$, c) and pro-inflammatory macrophages (LyC6$^{High}$, d) in regenerating QC muscles of Manf$^{fl/fl}$ and Manf$^{LysM\Delta}$ mice at 3dpi (n=4/condition). **e,** Representative plots of flow cytometry analysis of macrophage subpopulations, gated on the CD11b$^{pos}$ population, for single cell suspensions isolated from QC muscles of Manf$^{fl/fl}$ and Manf$^{LysM\Delta}$ mice at 2dpi, 0h and 16h after culture. Frequencies are from CD11b$^{pos}$ parent population. **f-i,** Ratio of Ly6C$^{Int}$ to Ly6C$^{High}$ macrophages (f,i), Ly6C$^{Low}$ to Ly6C$^{Int}$ macrophages (g); and percentage of myeloid cells (h), quantified by flow cytometry, in single cell suspensions isolated from QC muscles of Manf$^{fl/fl}$ and Manf$^{LysM\Delta}$ mice at 2dpi, 0h and 16h after culture, with or without recombinant MANF supplementation (f-h, n=4-5/condition; c, i=8/condition). Data are represented as average ± s.e.m. and each n represents one animal. p values are from two-tailed Student's t-test.

**a**

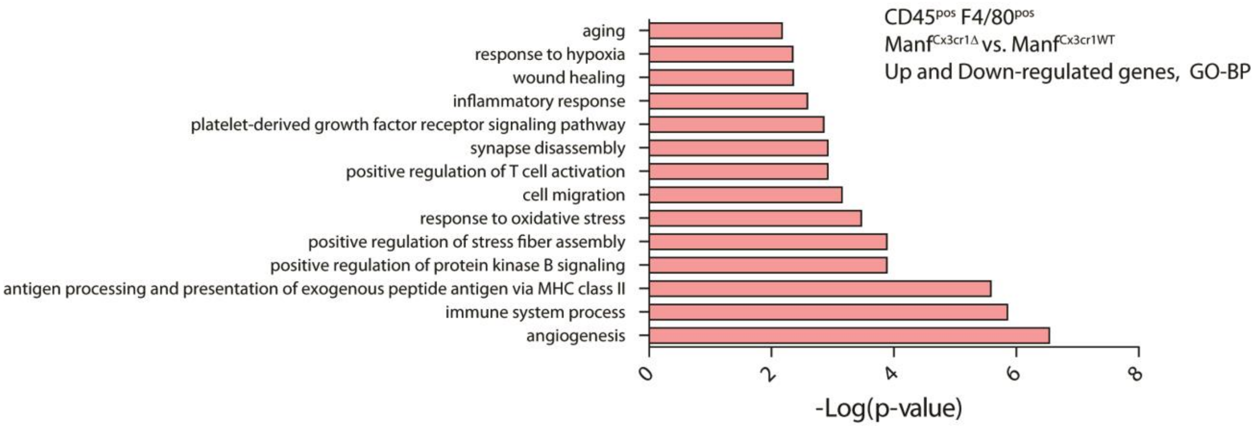


**b**

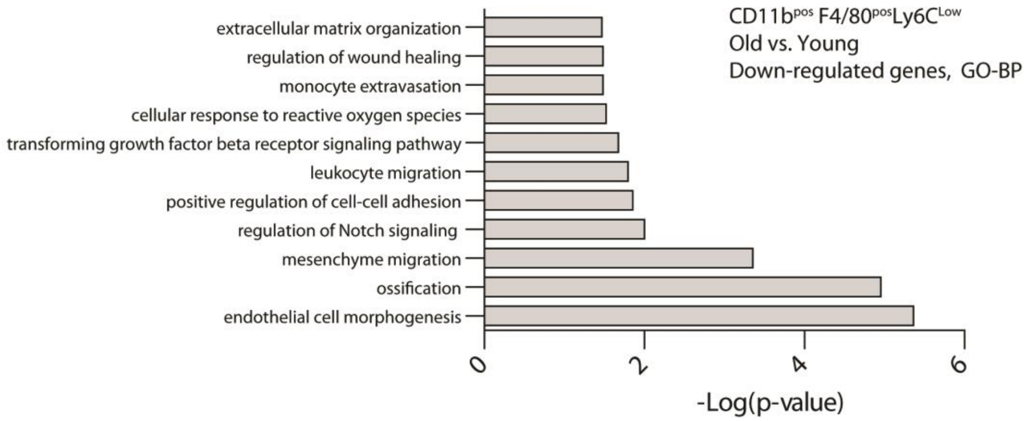


**Extended Data Figure 4. Defects of MANF-deficient and aged macrophages**

**a,** GO categories of biological processes showing significant enrichment in the dataset of genes differentially expressed in macrophages ($CD45^{pos}F4/80^{pos}$) FACS-isolated at 3dpi from QC muscles of $Manf^{Cx3cr1\Delta}$ mice compared to $Manf^{Cx3cr1WT}$ mice (fold change<0.75 or >1.5 and p≤0.05, p values from two-tailed Student's t-test, n=3/condition). **b,** GO categories of biological processes with relevance within the context of tissue regeneration showing significant enrichment in the dataset of genes down-regulated in pro-repair macrophages ($CD11b+F4/80^{pos}Ly6C^{Low}$) FACS-isolated at 3dpi from QC muscles of old (22-24mo) mice compared to yg (2-6mo) mice (fold change <0.75 and p≤0.05, p values from two-tailed Student's t-test, n=4/condition). GO, Gene Ontology; BP, Biological process.

**Supplementary Table 1**

Primer list for genotyping.

| Allele | Size (bp) | Forward Primer | Reverse Primer |
|---|---|---|---|
| **MANF Wt** | 512 | TGAAGCAAGAGGCAAAGAGAATCGG | TGCTCAGCTGCAGAGTTAGAGTTCC |
| **MANF fl** | 718 | TGAAGCAAGAGGCAAAGAGAATCGG | TGCTCAGCTGCAGAGTTAGAGTTCC |
| **Cx3Cr1 wt** | 695 | AAGACTCACGTGGACCTGCT | CGGTTATTCAACTTGCACCA |
| **Cx3Cr1 Cre-ER** | 300 | AGGATGTTGACTTCCGAGTTG | CGGTTATTCAACTTGCACCA |
| **R26 wt** | 198 | CTG GCT TCT GAG GAC CG | CCG AAA ATC TGT GGG AAG TC |
| **R26 Cre-ER** | 150 | CGT GAT CTG CAA CTC CAG TC | AGG CAA ATT TTG GTG TAC GG |
| **LyzM WT** | 350 | TTA CAG TCG GCC AGG CTG AC | CTT GGG CTG CCA GAA TTT CTC |
| **LyzM Cre** | **700** | CCC AGA AAT GCC AGA TTA CG | |

**Supplementary Table 2**

Primary Antibodies for IHC

| Target Gene | Species | Source | Dilution |
|---|---|---|---|
| MANF | rabbit | Sigma (SAB3500384) | 1:300 |
| F4/80 | rat | BioRad (MCA497G) | 1:40 |
| eMHC | mouse | D.S.H.B (F1.652) | Non-diluted |

**Supplementary Table 3**

Fluorophore-conjugated antibodies for FC analysis and FACS

| Fluorophore | Antibody | Source | Dilution |
|---|---|---|---|
| FITC | Anti-mouse CD45 | Biolegend (103107) | 1:200 |
| PE | Anti-mouse F4/80 | Biolegend (123110) | 1:50 |
| APC | Anti-mouse Ly6C | eBioscience™ (17-5932-82) | 1:200 |
| FITC | Anti-mouse Ly6G | Biolegend (127606) | 1:400 |
| APC-eFluor®780 | Anti-mouse CD11b | eBioscience™ (47-0112-82) | 1:200 |
| Alexa Fluor488 | Anti-mouse CD31 | Biolegend (102513) | 1:50 |
| PE-Cy5 | Anti-mouse CD45 | Biolegend (103109) | 1:100 |
| PE-Cy7 | Anti-mouse Sca-1 | Biolegend (108113) | 1:100 |
| PE | Anti-mouse α7integrin | Miltenyi Biotec (130-120-812) | 1:40 |

**Supplementary Table 4**

Primary antibodies for WB

| Target Gene | Species | Source | Dilution in WB |
|---|---|---|---|
| MANF | rabbit | Sigma (SAB3500384) | 1:1000 |
| Beta-Actin | mouse | DSHB (JLA-20C) | 1:200 |
| Vinculin | mouse | Sigma (V9131) | 1:2000 |

**Supplementary Table 5**
Primer list for RT-qPCR

| Gene | Species | Forward Primer | Reverse Primer |
|---|---|---|---|
| MANF | mouse | GCTGCCACCAAGATCATCAA | CACAGGGATATGGTGGGCC |
| Beta-actin | mouse | GCTCTGGCTCCTAGCACCAT | GCCACCGATCCACACAGAGT |
